# Cell cycle regulation of mitochondrial protein import revealed by genome-scale pooled bimolecular fluorescence complementation screening

**DOI:** 10.1101/770669

**Authors:** Kim Blakely, Patricia Mero, Roland Arnold, Ayesha Saleem, Christine Misquitta, Dahlia Kasimer, Sachin Kumar, Andrea Uetrecht, Kevin A. Brown, Alessandro Datti, David Hood, Philip Kim, Jason Moffat

## Abstract

A central focus of systems biology is the functional mapping of protein-protein interactions under physiological conditions. Here we describe MaGiCaL-BiFC, a lentivirus-based bimolecular fluorescence protein-fragment complementation approach for the high-throughput, genome-scale identification of protein-protein interactions in mammalian cells. After developing and validating this methodology using known protein-protein interaction pairs, we constructed genome-scale pooled BiFC libraries using the human ORFeome cDNA collection. These pooled libraries, containing ∼ 12,000 unique human cDNAs, were used to screen for candidate interaction partners of the mitochondrial transmembrane protein TOMM22. Following infection of cells with the TOMM22 bait and the pooled cDNA libraries, cells harboring candidate TOMM22 interacting proteins were isolated from the cell pool via fluorescence activated cell sorting, and identified via microarray analysis. This approach identified several known interaction partners of TOMM22, as well as novel physical and functional partners that link the mitochondrial network to proteins involved in diverse cellular processes. Notably, protein kinase CK2 was identified as a novel physical interaction partner of human TOMM22. We found that this association occurs preferentially during mitosis and involves direct phosphorylation of TOMM22, an event that may lead to attenuation of mitochondrial protein import. Together, this data contributes to the growing body of evidence suggesting eloquent coordination between cell cycle progression and mitochondrial physiology. Importantly, through high-throughput screening and focused validation, our study demonstrates the power of the MaGiCaL-BiFC approach to uncover novel functional protein-protein interactions, including those involving proteins with membrane-spanning domains, or of a transient nature, all within their native cellular environment.

## INTRODUCTION

Currently, new protein-protein interactions (PPIs) are most commonly identified using yeast two-hybrid (Y2H) methods (Fields and Song 1989; Rual *et al*. 2005) or affinity purification followed by mass spectrometry (AP-MS) (Ewing *et al*. 2007; Mak *et al*. 2010). While such techniques have offered great insight into the PPI networks in some mammalian cell types, there are disadvantages inherent to these methodologies. Notably, both technologies face severe limitations when trying to examine the interaction profiles of membrane-bound proteins. In recent years, development of the protein fragment complementation assay (PCA), including the biomolecular fluorescence complementation (BiFC) approach, have reformed the study of protein-protein interactions *in vivo* (Hu *et al*. 2002; Remy and Michnick 2004; Kerppola 2008b). These assays are based on the observation that the two halves of a rationally dissected protein can reconstitute *in vivo* to create a functional protein. Such technologies have been applied with success in mammalian cells using transfection-based methodologies (Hu *et al*. 2002; Remy and Michnick 2004), as well as an enhanced retroviral mutagen technique (Ding *et al*. 2006). Despite this, application of these technologies remains limited as transfection-based techniques are constrained to use in transfection-amenable cell types, and are often subject to high false positive rates, while the retroviral method faces limitations in terms of applicability to a wide range of mammalian cell types and the lack of full-length transcripts generated using this technology. Although no technique is likely to provide a perfect solution to these issues, an *in vivo* mammalian-based system bridging recent advances in both pooled-barcoded library screening and lentiviral-based technologies would help provide a solution to a number of these issues. Lentiviral expression systems encoding in-frame fusions to full-length human genes can be easily introduced into virtually any mammalian cell type, and expression levels off lentiviral vectors are more easily controlled compared to transfection-based techniques.

To this end, we have designed MaGiCaL-BiFC: a <u>Ma</u>mmalian, <u>G</u>ateway-<u>C</u>ompatible (Walhout *et al*. 2000), <u>L</u>entiviral-based (Zufferey *et al*. 1997; Dull *et al*. 1998; Moffat *et al*. 2006), <u>Bi</u>molecular <u>F</u>luorescence protein-fragment <u>C</u>omplementation assay (Michnick *et al*. 2000; Remy *et al*. 2002; Michnick 2003; Remy *et al*. 2007a; Remy *et al*. 2007b; Remy and Michnick 2007; Kerppola 2009; Schutze *et al*. 2009) based on the reconstitution of a split fluorescent protein, Venus (Nagai *et al*. 2002), for the genome-scale analysis of PPIs in mammalian cells. In MaGiCaL-BiFC, the two fragments of Venus are tagged to proteins of interest using Gateway recombinant technology (Hartley *et al*. 2000), packaged into lentivirus, and stably expressed as fusions within cells. When expressed separately, the split Venus fragments do not form a functional fluorophore (Kerppola 2008a). However, when each fragment is fused to one protein of an interacting pair, Venus is reconstituted creating a stable and quasi-irreversible fluorescent complex (Lalonde *et al*. 2008). The advantages of MaGiCaL-BiFC over currently used methodologies to study PPIs in mammalian cells are numerous: PPIs are detected *in vivo*, occur in their native cellular compartment, and most importantly, their localization can be visualized via microscopy, providing powerful insight into the biological function of specific PPIs. The lentiviral nature of this system allows for pooled genome-wide screening of PPIs in any cell line of interest using fluorescence activated cell sorting (FACS) (Mak *et al*. 2011). In addition, the pooled genetic screening approach involves minimal cost and is compatible with sequencing or microarray de-convolution.

## MATERIALS AND METHODS

### Cell lines

HEK293T cells were cultured in Dulbecco’s modified Eagle’s medium (DMEM) supplemented with 10% heat-inactivated fetal bovine serum (iFBS) and 1% penicillin/streptomycin (pen-strep) (Wisent, St. Bruno, QC). HeLa cells were cultured in McCoy’s 5A medium supplemented with 10% fetal bovine serum (FBS) and 1% pen-strep. The ReNcell VM NSC line was obtained from Millipore (Billerica, MA) and cultured according to the manufacturer’s protocol. Fibroblast growth factor (FGF) and epidermal growth factor (EGF) were purchased from BD Biosciences (Bedford, MA).

### Antibodies

Primary antibodies against TOMM22 (ab57523), TOMM20 (ab56783), CSNK2A1 (ab70774), MYC-HRP (ab1326) and FLAG-HRP (ab49763) were from Abcam (Cambridge, MA). HRP-conjugated secondary antibodies for Western blot detection were from Cell Signaling Technologies (NEB; Ipswich, MA). Primary antibodies against cyclins (sampler kit 9869) and for co-IP detection of CSNK2A1 (2656) were from Cell Signaling Technologies.

### Construction of the MaGiCaL-BiFC destinations vectors

The MaGiCaL-BiFC destination vectors were generated by PCR amplification of the CMV promoter and Gateway cassette with flanking N- and C-terminal Venus fragments from the vectors RFB-VF1, RFB-VF2, VF1-RFA and VF2-RFA (generous gift of Dr. Stephen Michnick) and cloned into pLJM1 (Sancak *et al*. 2008) using *Sna*BI and *Bst*BI restriction sites. The bait vectors were further modified by the addition of an IRES-RFP-WPRE fragment cloned between the *Xcm*I and *Nsi*I sites to replace the PGK promoter and PAC resistance gene.

### Proof-of-concept localization studies

Full-length cDNAs were obtained from the human ORFeome v5.1 entry clone collection (Thermo Fisher, Open Biosystems) and sequence verified. Gateway LR reactions were performed according to the manufacturer’s protocol (Life Technologies, Carlsbad, CA) to introduce the cDNA inserts into the MaGiCaL-BiFC destination vectors. Lentivirus was used to infect HEK293T and ReNcell VM NSC cultures at an MOI of ∼ 1. Four-days post-infection the cells were imaged via confocal microscopy. For NSC differentiation, bFGF and EGF were removed from the growth media four-days post-infection and cells were allowed to differentiate for one week. Media was changed every two days.

### Construction of the MaGiCaL-BiFC hORFeome v5.1 libraries

The hORFeome v5.1 entry clone library was pooled into 43 sub-pools of ∼ 370 clones each, and each pool was standardized to a concentration of 150 ng/µL. A standard LR reaction using the prey vectors, pLD-CMVpr-VF2-Gateway-hPGKpr-puro and pLD-CMVpr-Gateway-VF2-hPGKpr-puro, was performed with each pool for a total of 86 LR reactions. Following proteinase K inactivation, each reaction was transformed into ElectroMAX DH5-alpha_E cells (Life Technologies) and plated on LB agar + 100 µg/mL ampicillin. Overnight incubation at 37°C generated ∼ 400,000 colonies per transformation, for a ∼ 1000-fold representation of each pool. Colonies from each individual plate were pooled, and plasmid DNA extracted using a QIAfilter Plasmid Maxi Prep kit (Qiagen, Valencia, CA). DNA was standardized to 500 ng/µL, and pooled together to generate separate N- and C-terminal MaGiCaL-BiFC libraries.

### Lentivirus production

For the VF1-TOMM22 and prey library virus, ten million cells were seeded into 150 mm dishes twenty-four hours prior to transfection in DMEM supplemented with 10% iFBS and 0.1% pen-strep. The bait or prey-library plasmid DNA (18 μg), along with the packing plasmid psPAX2 (16.2 μg) and envelope plasmid pMD2.G (1.8 μg) were transfected using Fugene 6 (Roche, Basel, Switzerland) according to the manufacturer’s protocol. Eighteen hours post-transfection media was replaced with DMEM containing 1.1% bovine serum albumin (BSA) and 1% pen-strep. Virus was harvested at 42 h and 66 h post-transfection, pooled, aliquoted and stored at −80°C. Small-scale virus preparations were prepared in 60 mm plates using area-adjusted reagents.

### The MaGiCaL-BiFC pooled-screening procedure

HEK293T cells were co-transduced with VF1-TOMM22 and either the NVF2- or CVF2-prey library virus at a MOI of ∼ 0.5 each, such that ∼ 25% of cells contained both a bait and prey construct. Twenty-four hours post-infection the cells were selected with 2 µg/mL puromycin (Wisent). Four-days post-infection the cells were washed once with warm PBS, and collected by trypsinization. Cells were resuspended in PBS/2% iFBS containing 2 mM EDTA and counted.

One million cells were pelleted and frozen at −80°C as a pre-sort background sample, and the remaining cells were analyzed on a BD Facsaria, sorting for RFP^+^Venus^+^ events. For the N-terminal library screens, approximately 5 million cells were sorted of which 27.36% +/-3.78% were RFP+ and 0.079% +/-0.042% were RFP+/Venus+, providing a ∼ 125-fold coverage of the library ORFs screened. For the C-terminal library screens, approximately 3 million cells were sorted of which 25.89% +/-1.26% were RFP+ and 0.069% +/-0.012% were RFP+/Venus+, providing a ∼ 75-fold coverage of the library ORFs screened (Table S1 and Figure S4). Sorted cells were imaged post-sort on a WaveFX confocal spinning disc microscope (Quorum Technologies) using standard filter sets for FITC (Venus) and TexasRed (RFP) to confirm Venus expression in the sorted cells. Sorted cells were expanded for one week in culture before harvesting.

### Identification of positive hits from sorted populations

Genomic DNA was isolated from the expanded cells, and unsorted control populations, using the DNeasy Blood and Tissue Kit (Qiagen). The cDNA inserts were PCR-amplified using primers specific for either the N- or C-terminal libraries using Phusion DNA polymerase (New England Biolabs, Ipswich, MA). Amplicons were PCR-purified using the QIAquick PCR Purification Kit (Qiagen), and subsequently labeled using the BioPrime DNA Labeling System (Invitrogen). Biotinylated probes were hybridized to GeneChip Human Gene 1.0 ST Arrays (Affymetrix, Santa Clara, CA) according to the manufacturer’s protocol. Chip files were analyzed using Affymetrix Expression Console Software using hORFeome5-1 analysis file and RMA-Sketch normalization for analysis, with results in log form. The 3 replicates of the N-terminal dataset were analyzed together, and the background chip was analyzed separately. Similar analysis was done for C-terminal sets. The resulting files were then exported as .txt files using the annotation merge file hORFeome.hugene-1_0-st.v1_na28_hg18. The microarray includes 15,347 probesets that measure expression levels of 12,556 Entrez Gene IDs. For each of the screens, array expression values were floored at 1, and probesets were filtered to remove those with high average background from duplication measurements (bg > 1000) or low median expression from triplicate experiments (expr < 200). For the remaining probesets, the ratio of the expression intensity to background intensity was computed, along with the median and median absolute deviation (MAD) of the ratios. Hits were defined as probesets with ratios that were more than three MADs above the median value. For N-terminal screens, this threshold was 2.24, while for the C-terminal screens the threshold was 1.73. There was 294 probesets representing 263 unique Entrez Gene IDs called hits in the N-terminal screen, while there was 152 probesets representing 130 unique Entrez Gene IDs in the C-terminal screen (Table S2).

### Pair-wise BiFC validation

Full-length cDNA clones for select putative interaction partners of TOMM22 were picked from the human ORFeome v5.1 collection and sequence validated. Clones were transferred into the MaGiCaL-BiFC destination vectors by LR site-specific recombination, and used to generate lentivirus. HEK293T cells were co-transduced with VF1-TOMM22 and the select baits at a MOI of 1-5. Four days post-infection, cells were imaged on a WaveFX spinning disc confocal microscope using standard filter sets for FITC (Venus) and TexasRed (RFP).

### Secondary siRNA screen

The siRNA secondary screen was performed by transfecting HEK293T cells seeded in 384-well plates with Dharmacon siRNA SMARTpools, with each pool containing 4 siRNAs against a given target. Each transfection was performed in duplicate. Three days post-transfection, cells were stained with MitoTracker Green FM, MitoTracker Red CMXRos, and Hoechst 33342. Plates were imaged using a GE Healthcare IN Cell Analyzer 2000 epifluorescent HTS microscope. Images were acquired using a 10X long working-distance 0.45NA objective (Nikon) and standard filter sets for FITC (MitoTracker Green FM), Cy3 (MitoTracker Red CMXRos) and DAPI (Hoechst). Nine fields of view per well were captured. Images were analyzed using Perkin Elmer’s Columbus image analysis server running Acapella 2.0. Reported values represent the average from duplicate wells.

### Protein extraction and immunoblotting

Cells were lysed with RIPA lysis buffer containing protease inhibitor cocktail (Sigma) and quantified by BCA (Pierce). Lysates (10-40 μg/lane) were separated by SDS-PAGE and protein was transferred onto a polyvinylidene fluoride (PVDF) membrane (Hybond-P, GE Healthcare, Uppsala, Sweden). Specific primary antibodies (see Antibodies subsection) were detected using the appropriate secondary HRP-conjugated antibodies and visualized by enhanced chemiluminescence detection (Supersignal West Pico; Pierce - Thermo Fisher Scientific).

### Exogenous co-immunoprecipitation experiments

Full-length cDNA clones for TOMM34 and CSNK2A1 were picked from the human ORFeome v5.1 collection and sequence verified. Clones were transferred into a N-terminal MYC-tagged lentiviral destination vector by LR site-specific recombination. N-terminal FLAG-tagged TOMM22 was generated by LR site-specific recombination of the TOMM22 entry clone into the MAPLE lentiviral destination vector (Mak *et al*. 2010). HEK293T cells were co-transfected with TOMM22 and the select baits, and were subsequently immunoprecipitated using αFLAG M2-conjugated agarose beads (Sigma-Aldrich, St. Louis, MO). Bound proteins were resolved by SDS-PAGE and detected using an αMYC-tag HRP-conjugated antibody (Abcam).

### Cell cycle blocks

HeLa cells were seeded in 150 mm plates in normal growth media at a density of 2.5 million cells/plate. Twenty-four hours post-seeding, media was exchanged for media containing 4 mM thymidine (Sigma-Aldrich). Following an overnight incubation (16 h), cells were released into fresh media for 7.5 h. Media was again exchanged for media containing 4 mM thymidine or 200 ng/mL nocodazole (Sigma-Aldrich). Following an overnight incubation (16 h), cells were released into fresh media for varying amounts of time. Release of double thymidine-blocked cells for 0, 2.5, 5.5 and 7.5 h generated synchronous cell populations at approximately G1-S, early S, late S and S-G2 phases of the cell cycle. Release of thymidine/nocodazole blocked cells for 0.5 and 3 h generated synchronous cell populations at approximately M and G1 phases of the cell cycle. Cells were collected to prepare lysates for Western blot analysis and confirmation of the cell cycle phases using specific primary antibodies against Cyclin A, Cyclin B1, Cyclin E2 or Cyclin E (Cell Signaling Technology), or for endogenous co-IPs.

### Endogenous IP of TOMM22 and protein kinase CK2 in asynchronous and M-phase blocked cells

Asynchronously growing HeLa cells and HeLa cells blocked in M-phase were harvested in co-IP buffer (50 mM Hepes-KOH, 0.1% Triton-X100, 2 mM EDTA, 100 mM NaCl, and 1X protease inhibitors (Sigma-Aldrich)) and lysed for 30 minutes on ice. Lysates were cleared by centrifugation at 14,000 g for 10 minutes at 4°C. Lysates were quantified by BCA and equal amounts of asynchronous or M-phase lysates were used for the IP of endogenous TOMM22 or CSNK2A1 using specific primary antibodies (see Antibodies subsection) overnight at 4°C with rotation. The next day protein G Ultralink beads (Pierce-Thermo Fisher Scientific) were added to the samples, which continued to rotate for an additional 2 h at 4°C. Beads were collected by centrifugation at 2,200 RPM, and washed 3 times with ice cold PBS. Bound proteins were resolved by SDS-PAGE, and used for subsequent Western blotting for detection of immunoprecipitated TOMM22 and CSNK2A1.

### Generation of phosphorylation site mutations in TOMM22

Mutagenesis primers were used to introduce phosphorylation site mutations into the TOMM22 entry clone using the GeneArt Site-Directed Mutagenesis System (Life Technologies). Primers specific to serine residue 15 and threonine residue 43 were each individually mutated to alanine (A) and glutamic acid (E). In addition, the double alanine (AA) and double glutamic acid (EE) mutants were generated. Common primers were used to amplify these ORFs to introduce a single N-terminal FLAG tag into the construct, as well as an in-frame stop codon. These entry clones were then introduced into the pMAL vector (generous gift of L. Naldini) by site-specific LR reaction, and virus was generated as described above. HEK293T cells were transduced with the lentiviral constructs, or a control *Renilla luciferase* (RLUC) construct designed with an N-terminal FLAG tag. Two-days post infection, cells were infected with an shRNA targeting the 3’UTR of TOMM22 to deplete the endogenous TOMM22 transcript.

### Mitochondrial isolations

Mitochondrial isolations were performed as described previously (Frezza *et al*. 2007). Briefly, five 150 mm tissue-culture dishes were seeded with 10 million cells for each of the mutant cell lines two days prior to the experiment. On the day of isolation, cells (∼ 200 million per mutant) were washed twice with ice-cold PBS, and collected using a cell scraper. Cells were pelleted at 600 g at 4°C for 10 minutes, resuspended in 3 mL ice-cold IBC buffer (200 mM sucrose, 10 mM Tris-MOPS pH 7.4, 100 nM EGTA/Tris) and homogenized using a Teflon pestle/glass potter operated at 1500 RPM for 30 strokes. Nuclei were pelleted by centrifugation at 600 g at 4°C for 10 minutes. The supernatant was collected and pelleted at 7000 g at 4°C for 10 minutes. The mitochondrial pellet was washed once with ice-cold IBC buffer, and a fraction was taken for lysis in RIPA buffer, followed by quantification by BCA.

### *In vitro* transcription, translation and import of OCT

The *in vitro* mitochondrial import assays were performed as described previously (Singh and Hood 2011). Briefly, vector encoding full-length ornithine transcarbamylase (OCT) was linearized using *Sac*I and subsequently phenol extracted and ethanol precipitated. OCT was *in vitro* transcribed at 40°C for 90 min using SP6 RNA polymerase, followed by *in vitro* translation in the presence of [^35^S]methionine within a rabbit reticulocyte lysate system. For the mitochondrial protein import assay, 50μg of mitochondria were preincubated for 10 min at 30°C before the import assay. Translation mix (12 μl) was added to the mitochondria, and the import incubation was allowed to proceed at 30°C for 8 min. Subsequently, import was halted by the addition of an aliquot of the mitochondrial translation mix to an ice-cold sucrose cushion (600 mM sucrose, 100 mM KCl, 20 mM HEPES, and 2 mM MgCl2). Mitochondria were pelleted by centrifugation for 15 min at 16,000 g at 4°C and resuspended in 20 μl of ice-cold breaking buffer (600 mM sorbitol, 20 mM HEPES, pH 7.4). The samples were then denatured at 95°C in the presence of lysis buffer for 5 min, quick cooled on ice, and then resolved by SDS-PAGE (12% gel). Gels were processed and dried as described elsewhere (Craig *et al*. 1998). Images and subsequent quantification were obtained with electronic autoradiography (Quantity One, Bio-Rad, Hercules, CA). Imported mature OCT (mOCT) was distinguished from precursor OCT (pOCT) by its lower molecular weight. The percentage of imported protein was calculated based on the ratio of the intensity of the mOCT to total OCT protein (mOCT+pOCT). A Repeated Measures ANOVA was used to evaluate the significance of the difference between the A and E mutants. Each mutant was evaluated in three biologically independent experiments, following fresh lentiviral infections and mitochondrial isolations.

## RESULTS

### Design of the MaGiCaL-BiFC system

We designed MaGiCaL-BiFC to enable genome-wide analyses of PPIs in any mammalian cell line of interest using any bait protein of interest, based on the bimolecular fluorescence protein-fragment complementation assay and lentiviral-based pooled barcode screening methodology. To achieve this, a series of lentiviral (Zufferey *et al*. 1997; Dull *et al*. 1998) destination vectors were developed that incorporate a Gateway expression cassette (Walhout *et al*. 2000) in frame with either the N- or C-terminal fragment of the rationally-dissected fluorescent protein Venus (VF1 and VF2, respectively) (Nagai *et al*. 2002; MacDonald *et al*. 2006). To circumvent issues that may arise due to orientation of the tag, vectors were built in both orientations, such that cDNAs could be tagged on either terminus. The bait vectors (containing VF1) include an internal ribosome entry site (Van Der Kelen *et al*. 2009) in frame with red fluorescent protein (IRES-RFP) to gauge bait expression levels. Prey vectors (containing VF2) include the puromycin resistance gene puromycin N-acetyltransferase (PAC) under control of the constitutive human phosphoglycerate kinase promoter (hPGKpr-puro) to allow efficient selection of prey-expressing cells (Figure 1A).

**Figure 1.**
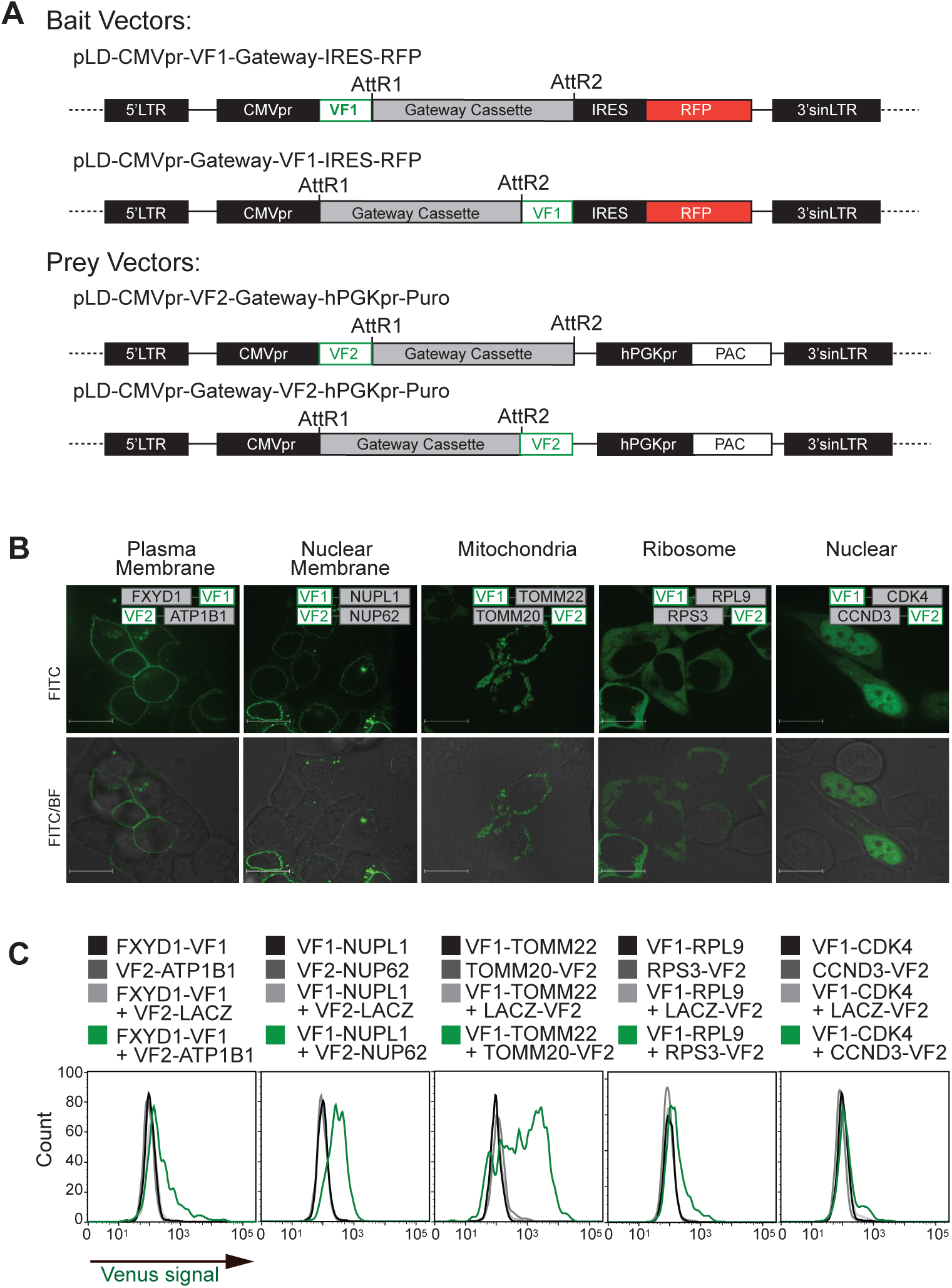
MaGiCaL-BiFC system. (A) Schematic of the MaGiCaL-BiFC lentiviral destination vectors containing a Gateway expression cassette in-frame with either the N- or C-terminal fragment of the fluorescent protein Venus (VF1 and VF2, respectively). The bait vectors include an internal ribosome entry site in frame with the gene encoding the red fluorescent protein (IRES-RFP). Prey vectors include a puromycin resistance gene (PAC) under the control of the constitutive human phosphoglycerate kinase promoter (hPGKpr-PURO). (B) Series of MaGiCaL-BiFC assays in different cellular compartments. Expression clones corresponding to protein-protein interaction pairs with well established cellular localization patterns were generated for the plasma membrane, FXYD domain containing ion transport regulator 1 (FXYD1) and ATPase, Na+/K+ transporting, beta 1 polypeptide (ATP1B1); ribosome, ribosomal protein L9 (RPL9) and ribosomal protein S3 (RPS3); mitochondria, translocase of outer mitochondrial membrane 22 homolog (yeast) (TOMM22) and translocase of outer mitochondrial membrane 20 homolog (yeast) (TOMM20); nuclear membrane, nucleoporin like 1 (NUPL1) and nucleoporin 62kDa (NUP62); and nucleus, cyclin-dependent kinase 4 (CDK4) and cyclin D3 (CCND3). Four-days post infection at an MOI<0.3, cells were imaged for Venus expression. Scale bar denotes 16 µm. (C) HEK293T cells infected (MOI<1) with the indicated lentiviruses were harvested and analysed by flow cytometry for Venus expression.

To test the sensitivity and accuracy of MaGiCaL-BiFC, we generated expression clones for established protein interactions with specific localization patterns (ie. “test set”) including FXDY1-ATP1B1 (plasma membrane), NUPL1-NUP62 (nuclear membrane), TOMM22-TOMM20 (mitochondria), CYP1A1-CYB5A (endoplasmic reticulum), RPL9-RPS3 (ribosome/cytoplasmic) and CDK4-CCND3 (nuclear). Vectors were generated by Gateway site-specific recombination, with one protein of the pair introduced into both orientations of the bait vector, and the other into both orientations of the prey vector. Lentiviruses from these expression constructs were generated and used to infect HEK293T cells either singly, as pairs (testing all four possible combinations) or with a negative control construct that includes beta-galactosidase (LACZ). Venus signal was undetectable by fluorescence microscopy when each bait-VF1 or prey-VF2 construct was expressed alone, and was extremely faint and/or uneven with the complementary negative control construct. When the correct pairs were expressed together, predicted signal was readily detectable by either microscopy (Figure 1B and Figure S1) or flow cytometry (Figure 1C).

To further assess our lentiviral system in cell types that are difficult to transfect, we infected human neural stem cells (ie. ReNcell VM cells), with the lentivirus test assays described above, and found the predicted Venus localization patterns (Figure S2A). Moreover, these stem cells were differentiated into neural cells by removing mitogens (bFGF and EGF) for one week, and then re-imaged (Figure S2B). Signal intensity and specificity were maintained in the differentiated neural cells, addressing the applicability of this system to cell types refractory to transfection-based PPI detection methods such as neural stem cells.

### Construction of the MaGiCaL-BiFC libraries

We next wanted to develop prey libraries to enable pooled genome-scale PPI screens by virtue of the fact that reconstituted Venus signal can be efficiently detected and sorted by FACS for many different protein-protein interaction assays (Figure 1C). To generate the N- and C-terminal VF2 lentiviral pooled libraries, the human ORFeome v5.1 entry clone library was pooled and introduced into the prey destination vectors by Gateway site-specific recombination to generate individual lentiviral-based expression plasmid pools with VF2 sequence in frame with either the N- or C-terminus of >12,000 unique human open reading frames (ORFs) (Figure S3A). Since each human ORF can act as a barcode, a long-range PCR procedure was developed with common amplification primers to generate labeled probe from the expression plasmid pools, as well as genomic DNA from HEK293T cells infected with the pooled lentivirus libraries in order to assess representation and quality of the ORFs post-infection. We estimated library representation to be >70% in the lentiviral plasmid pools and following packaging and infection into HEK293T cells (Figure S3B).

### Proof-of-concept screen for TOMM22 binding partners

We chose to investigate the PPI network of TOMM22, which plays a central role in cell metabolism, survival, and growth (Yano *et al*. 2000; Bellot *et al*. 2007). TOMM22 was an attractive target since it forms a complex with several additional components of the TOM complex, mediating protein translocation into mitochondria. Due to the orientation of the TOMM proteins in the outer mitochondrial membrane (OMM) (Figure 2A), only specific tagging orientations should generate positive fluorescent signal when these interactions are assessed by MaGiCaL-BiFC, as was observed in the pairwise tests described above (Figure 1B), providing an ideal scheme to test the methodology.

**Figure 2.**
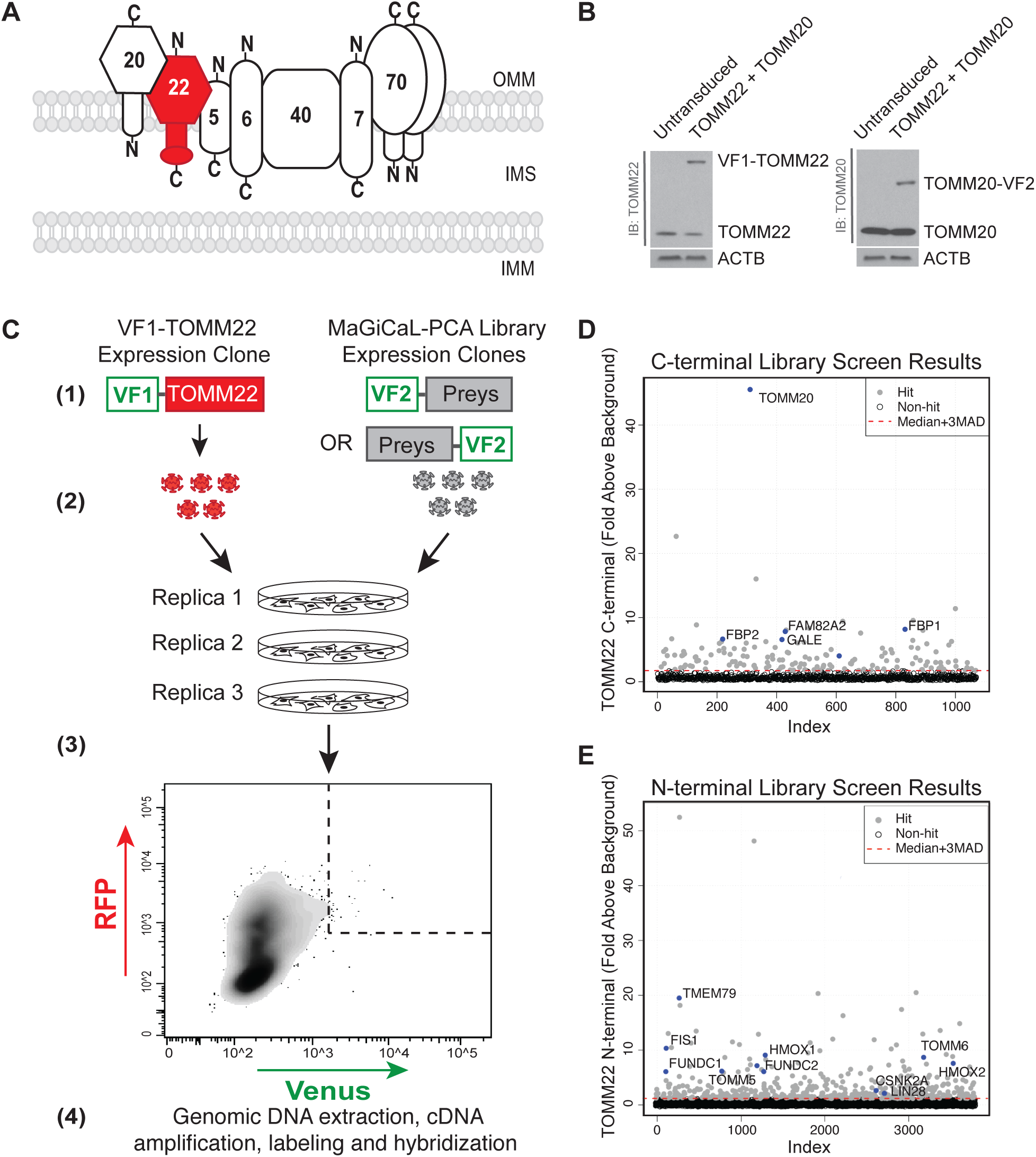
Proof-of-concept screen with TOMM22. (A) Schematic of the mitochondrial TOM complex showing orientation of the individual components at the outer mitochondrial membrane. (B) Western blot analysis for tagged and endogenous TOMM22 and TOMM20 in doubly infected HEK293T cells. (C) Schematic of the MaGiCaL-BiFC screening protocol, *(1)* MaGiCaL-BiFC expression clones were packaged as lentivirus particles, *(2)* lentivirus was collected and used to doubly infect HEK293T cells in triplicate at a MOI of ∼ 0.5 each, *(3)* four-days post infection RFP^+^Venus^+^ cells were collected, *(4)* genomic DNA was extracted from expanded cells for PCR amplification of the cDNA inserts. Following amplicon labeling, samples were hybridized to microarrays to identify putative interactors. The results of the average of the 3 microarray hybridization experiments plotted as fold-enrichment over background as a function of probeset/index for the N-terminal screens (D) and the C-terminal screens (E).

To screen for TOMM22 interacting proteins with pools of VF2-tagged preys using cell sorting and microarray de-convolution, the VF1-TOMM22 construct, which has the VF1-tag facing the cytoplasm, was generated for use as the bait (Figure 2A). To ensure that the protein expression off the lentiviral constructs was not exaggerated relative to endogenous levels, which could lead to high false positive rates, the protein levels of VF1-TOMM22, and its known interaction partner TOMM20-VF2 (described above), were analyzed by Western blot following transduction into HEK293T cells and found to be similar or lower than endogenous TOMM22 and TOMM20 (Figure 2B), consistent with what has been observed for similar lentiviral vector systems (Mak *et al*. 2010). For screening, HEK293T cells were simultaneously infected with the VF1-TOMM22 bait virus and either the N- or C-terminally tagged prey virus libraries at a multiplicity of infection of ∼ 0.5 for each of the bait and prey, so that an average of 25% of the cells expressed both the bait and a prey construct. For the N-terminal library screens, approximately 5 million cells were sorted of which 27.36% +/-3.78% were RFP+ and 0.079% +/- 0.042% were RFP+/Venus+, providing a ∼ 125-fold coverage of the library ORFs screened (Table S1 and Figure S4). For the C-terminal library screens, approximately 3 million cells were sorted of which 25.89% +/-1.26% were RFP+ and 0.069% +/-0.012% were RFP+/Venus+, providing a ∼ 75-fold coverage of the library ORFs screened (Table S1 and Figure S4). Genomic DNA was harvested from the sorted cells, as well as a control population of unsorted cells, in order to amplify the ORF inserts using common primers that flank the Gateway cassette. Following amplicon labeling, samples were hybridized to oligonucleotide microarrays and subsequently scanned to identify enriched signals that represent putative TOMM22 interactors (Figure 2C).

In order to identify putative TOMM22 interacting partners, we developed an enrichment score r, the median ratio of probeset intensities to background for each gene, where a hit was defined as r greater than 3 MADs above the median. This equated to 263 and 130 unique genes from the N-terminal and C-terminal library screens, respectively (Table S2). The analysis revealed a number of expected positive controls; for example TOMM20, a well-characterized TOMM22 interactor, was identified as the top hit in the C-terminal screen (r=45.5), consistent with its orientation in the OMM (Figure 2D). Conversely, TOMM5 (r=6.7) and TOMM6 (r=9.2) were only detected in the N-terminal screen, as expected (Figure 2E). In addition to identifying several TOM complex proteins such as TOMM20, TOMM5 and TOMM6, we also identified additional OMM proteins related to apoptosis (e.g. BAX, BCL2L1, BCL2L2 and BCL2L13), mitochondrial morphology (e.g. FIS1), cytoskeletal function (e.g. CORO1B, PFN1, TWF20) and metabolism (e.g. MGLL, FBP2, G6PD, CDS2, LDHB, CYB5B) (Table S2). Notably, the pro-apoptotic BCL2-family member BAX is a well-characterized interaction partner of TOMM22 (Bellot *et al*. 2007), and was specifically identified in the N-terminal MaGiCaL-BiFC screen, consistent with previous reports on its mitochondrial membrane orientation (Suzuki *et al*. 2000).

To confirm a subset of putative VF1-TOMM22 orientation-specific interactions from the primary screening data, we constructed a selection of prey constructs tagged with VF2 at the N- and C-termini and tested these in bimolecular fluorescence complementation assays in HEK293T cells using the MaGiCaL-BiFC vectors. Proteins for follow-up were selected based on interest and prior knowledge, and were chosen to include those with variable functions and subcellular localization patterns. For instance, mitochondrial import and mitochondrial morphology have been shown to be tightly coordinated processes, although the mechanisms regulating this are only poorly understood (Stojanovski *et al*. 2006). Given that FIS1, which is known to regulate mitochondrial morphology, was identified as a strong TOMM22 interactor, we chose to pursue this interaction in follow-up. Other mitochondrial proteins, including FUNDC1 and FUNDC2 were chosen based on the principle that members of this family are newly implicated in autophagy (Liu *et al*. 2012). Given that the TOMM machinery has been shown to function in this process (Bertolin *et al*. 2013; Yang and Yang 2013), we thought that these interactions might help to shed additional light on this important cellular process.

Similarly, crosstalk between mitochondria and endoplasmic reticulum is an emerging area of study that leaves much to be uncovered (English and Voeltz 2013; Kornmann 2013). For that reason, we chose to investigate the putative endoplasmic reticulum proteins identified as interaction partners of TOMM22, including both HMOX1 and HMOX2. Other putative interaction partners such as GALE, LIN28, and TMEM79 were chosen based on the fact that they have little to no known prior information linking them to TOMM22, or mitochondria, and novel interactions with TOMM22 may help to provide additional insight into their biological functions. In total, 17 genes were tagged in both orientations and 14/17 showed the predicted localization pattern (Figure 3A). As expected, the TOMM5 and TOMM6 preys showed a strong fluorescence signal in the mitochondria with the bait TOMM22 in an orientation-specific manner (Figure 3A). The remaining 12 preys that showed a strong Venus fluorescence signal included FIS1, FAM82A2, HMOX1, HOMX2, FBP1, FBP2, GALE, LIN28, FUNDC1, FUNDC2, TMEM79 and REEP6 (Figure 3A). The Venus signals also co-localized with the mitochondrial stain Mitotracker Red CMXRos (Figure 3B). Notably, the co-localization patterns varied in that some interactions showed strong OMM staining (e.g. HMOX1, HMOX2) while others showed punctate mitochondrial staining (e.g. FAM82A2/RMD3). Overall, these observations suggest that our primary TOMM22 sorting screen revealed a number of known and potentially novel protein interactions with a component of the TOM complex.

**Figure 3.**
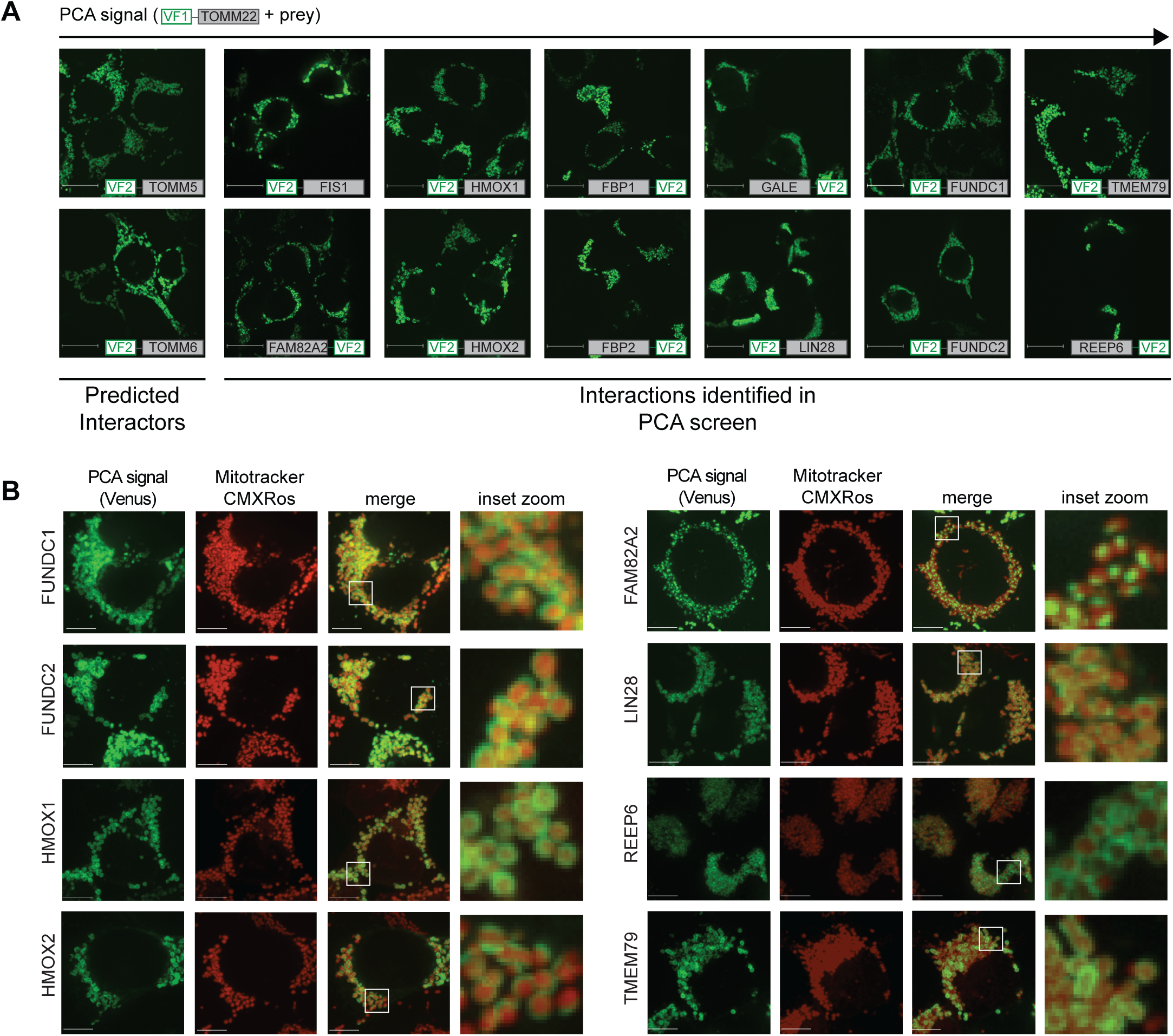
Confirmation of TOMM22 interactors by pairwise BiFC. (A) Confirmation of a subset of known and newly identified TOMM22 protein interactors. (B) Co-staining of TOMM22 protein interactions with Mitotracker to validate mitochondrial localization of the PCA signal.

### Link between TOMM22 interactors and mitochondrial mass/potential

We reasoned that some of the putative novel TOMM22 interactors that were identified in our screens could also be important for mitochondrial physiology. To test this hypothesis, we performed a secondary siRNA screen to determine the effects of knocking down the TOMM22 interaction partners on mitochondrial mass and/or potential. We chose 80 random putative interactors from both the N- and C-terminal screens and used high-content microscopy and quantitative image analyses to measure the changes in mitochondrial mass and/or potential following knockdown of these genes with siRNA pools and staining with Mitotracker dyes (Figure 4A). HEK293T cells were plated in 384-well plates and transfected in duplicate with siRNA pools targeting each of the 80 putative interaction partners. siRNAs targeting SIN3A, BAK, and MFN2, which have been previously reported to influence mitochondrial membrane potential, were included as controls. A scrambled siRNA pool that has no effect on membrane potential relative to untransfected cells was also included as a control. Three days post-transfection, the cells were stained with two different mitochondria-specific dyes, MitoTracker Green FM and MitoTracker Red CMXRos, and analyzed by high-content microscopy. Importantly, while both dyes accumulate in mitochondria based on mitochondrial mass, MitoTracker Red CMXRos is also sensitive to membrane potential. Thus, the red-to-green ratio can be used to evaluate changes in mitochondrial membrane potential specifically, as has been previously reported (Yoon *et al*. 2010).

**Figure 4.**
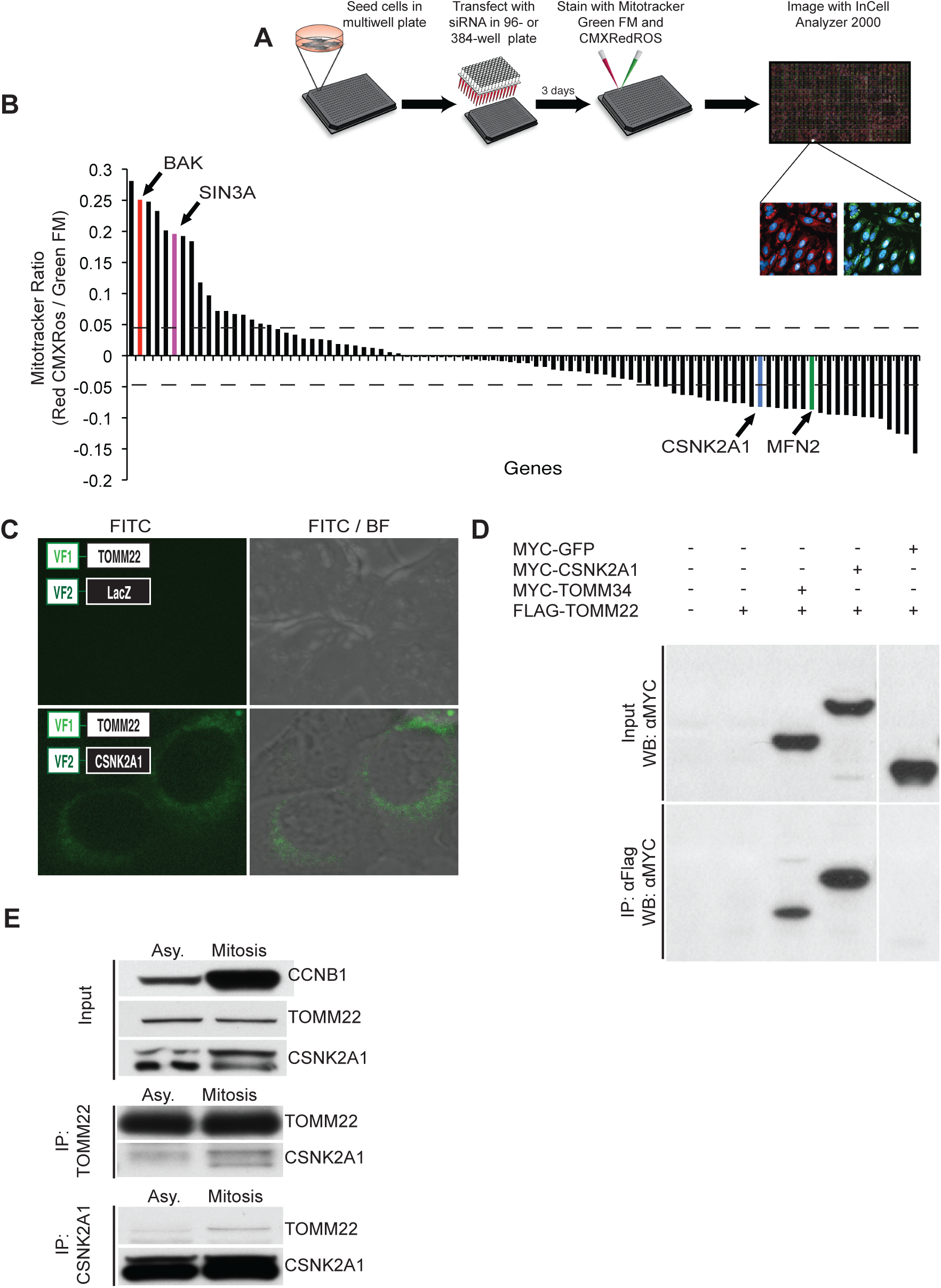
TOMM22 physically interacts with protein kinase CK2. (A) Schematic of secondary screen for effects of perturbing TOMM22 interactors on mitochondrial physiology. Cells transfected in 384-well plates with siRNA pools targeting select putative TOMM22 interaction partners were stained with MitoTracker Red CMXRos (which accumulates based on mass and potential) and MitoTracker Green FM (which accumulates based on mass). The ratio of the red-to-green signal is used as a relative measure of mitochondrial membrane potential. (B) Bar graph displays the genes ordered along the x-axis from highest to lowest relative membrane potential index (calculated relative to siScramble control wells). Dashed lines indicate 3 standard deviations above and below the siScramble membrane potential index. BAK (red bar), SIN3A (pink bar) MFN2 (green bar) and protein kinase CK2 (blue bar) are indicated. (C) MaGiCal-BiFC confirmation of the interaction between protein kinase CK2 and TOMM22. (D) Over-expression co-IP confirmation of the TOMM22-CSNK2A1 interaction. HEK293T cells were transfected with either N-terminally FLAG-tagged TOMM22 alone, or with N-terminally MYC-tagged TOMM34, protein kinase CK2 or GFP. TOMM22 was immunoprecipitated from cell lysates using αFLAG M2 agarose beads, and interactors were detected by Western blot using an αMYC-tag antibody. (E) Endogenous co-IP of protein kinase CK2 and TOMM22 from asynchronous and M-phase arrested cells.

Target genes were classified as hits if they altered the MitoTracker Red CMXRos-to-MitoTracker Green FM signal (reflecting a change in mitochondrial membrane potential) more than three standard deviations away from the mean of the siScramble control wells. The control gene, SIN3A, whose knockdown was previously reported to increase mitochondrial membrane potential (Yoon *et al*. 2010), came out among our top candidates that increased the MitoTracker Red CMXRos-to-MitoTracker Green FM signal ratio, indicating an increase in mitochondrial membrane potential (Figure 4B, *pink bar*). Also, the pro-apoptotic protein BAK came out among the top genes whose knockdown increased the MitoTracker Red CMXRos-to-MitoTracker Green FM ratio (Figure 4B, *red bar*), consistent with the pro-apoptotic role of BAK in mitochondrial outer membrane permeabilization and membrane potential dissipation (Breckenridge and Xue 2004). Conversely, knockdown of MFN2 decreased mitochondrial membrane potential (Figure 4B, *green bar*), which is consistent with previous reports (Chen *et al*. 2003). Using this approach, we found that siRNA-mediated knockdown of ∼ 57% of the 80 putative TOMM22 interaction partners identified in MaGiCaL-BiFC had an effect on mitochondrial membrane potential. This is in contrast to genome-scale approaches employing a similar method, which have reported that ∼ 3% of screened candidates affect membrane potential or mass when depleted (Yoon *et al*. 2010). This suggests that our MaGiCaL-BiFC hits are highly enriched for genes that impact mitochondrial physiology.

One of the candidates identified as a TOMM22 interaction partner, and whose knockdown in the secondary siRNA screen decreased mitochondrial membrane potential, was the protein kinase CK2 alpha (also known as CK2α or CSNK2A1), encoded by the CSNK2A1 gene (Figure 4B, *blue bar*). Interestingly, a physical and functional interaction was recently described between yeast CK2 and yeast Tom22 (Schmidt *et al*. 2011), which was reported to result in Tom22 phosphorylation, leading to increased localization of Tom22 to mitochondria, and ultimately increased protein import into mitochondria. Although human TOMM22 phosphorylation has not been studied in mammalian cells, phospho-TOMM22 peptides have been reported in large-scale phospho-proteomic screens (Dephoure *et al*. 2008; Olsen *et al*. 2010). Importantly, both of these phosphorylation sites, serine 15 and threonine 43, conform to the protein kinase CK2 phosphorylation consensus sequence, S/T-X-X-E/D, where protein kinase CK2 will phosphorylate a serine or threonine residue located N-terminally of acidic amino acid residues (Meggio and Pinna 2003). Moreover, both occur in the cytoplasmic domain of TOMM22. Interestingly, the previous detection of phospho-TOMM22 was reported in proteome-scale phosphoproteomic studies that analyzed the phosphoproteome under varying phases of the cell cycle. Both S15 (Olsen *et al*. 2010) and T43 (Dephoure *et al*. 2008) on TOMM22 were found to be phosphorylated specifically in cells synchronized at M-phase, suggesting that these are mitotic-specific phosphorylation events.

### Protein kinase CK2 interacts with and can phosphorylate TOMM22

To validate the interaction between TOMM22 and protein kinase CK2, CSNK2A1 was introduced into the N-terminally tagged MaGiCaL-BiFC prey vector, consistent with the tagging-orientation identified in the screen. VF1-TOMM22 and VF2-CSNK2A1 were co-transduced into HEK293T cells, and Venus signal was readily detectable by fluorescence microscopy compared to control infections, thus confirming the interaction between protein kinase CK2 and TOMM22 by this method (Figure 4C).

To validate the interaction between TOMM22 and protein kinase CK2 using an orthogonal protein interaction assay, co-immunoprecipitation (co-IP) experiments were performed. Briefly, TOMM22 was co-expressed in HEK293T cells as an N-terminally FLAG-tagged protein together with MYC fusion proteins corresponding to negative (GFP) and positive (TOMM34) control interaction partners, as well as protein kinase CK2. FLAG-tagged TOMM22 was immunoprecipitated (IP) using αFLAG M2-conjugated beads, and following washing, bound proteins were separated by SDS-PAGE and Western blots were performed using an anti-MYC antibody for detection of the co-immunoprecipitated interactors. No interaction could be detected between TOMM22 and GFP, however TOMM22 was effectively able to co-IP both TOMM34 and protein kinase CK2 (Figure 4D).

To further investigate the endogenous interaction between TOMM22 and protein kinase CK2, a series of cell cycle blocks using both thymidine and nocodazole to arrest HeLa cells in various phases of the cell cycle were performed, followed by IP of either TOMM22 or protein kinase CK2. Efficient block and release of the cells at the various cell cycle phases was assessed through Western blot analysis of the relative levels of various cyclin proteins (Figure S5). Importantly, we observed an interaction between TOMM22 and protein kinase CK2 with reciprocal IPs at endogenous protein levels (Figure 4E), which was more prevalent when using lysates from mitosis-arrested cells (Figure 4E), consistent with previous reports on TOMM22 phosphorylation status in human cell lines (Dephoure *et al*. 2008; Olsen *et al*. 2010). Taken together, these results suggest that protein kinase CK2 physically associates with TOMM22 in the cell, and this interaction is enhanced during mitosis.

### TOMM22 phosphorylation by protein kinase CK2 impacts mitochondrial protein import

We reasoned that because protein kinase CK2 associates with TOMM22, it might directly phosphorylate TOMM22. To confirm that protein kinase CK2 can directly phosphorylate TOMM22, *in vitro* phosphorylation assays were performed using recombinant GST-tagged TOMM22 and purified protein kinase CK2 holoenzyme in the presence of radiolabelled ATP. Protein kinase CK2 was able to effectively phosphorylate TOMM22 in a time-dependent manner (Figure 5A), suggesting that there may be a functional role for the protein kinase CK2-TOMM22 interaction in mammalian cells.

**Figure 5.**
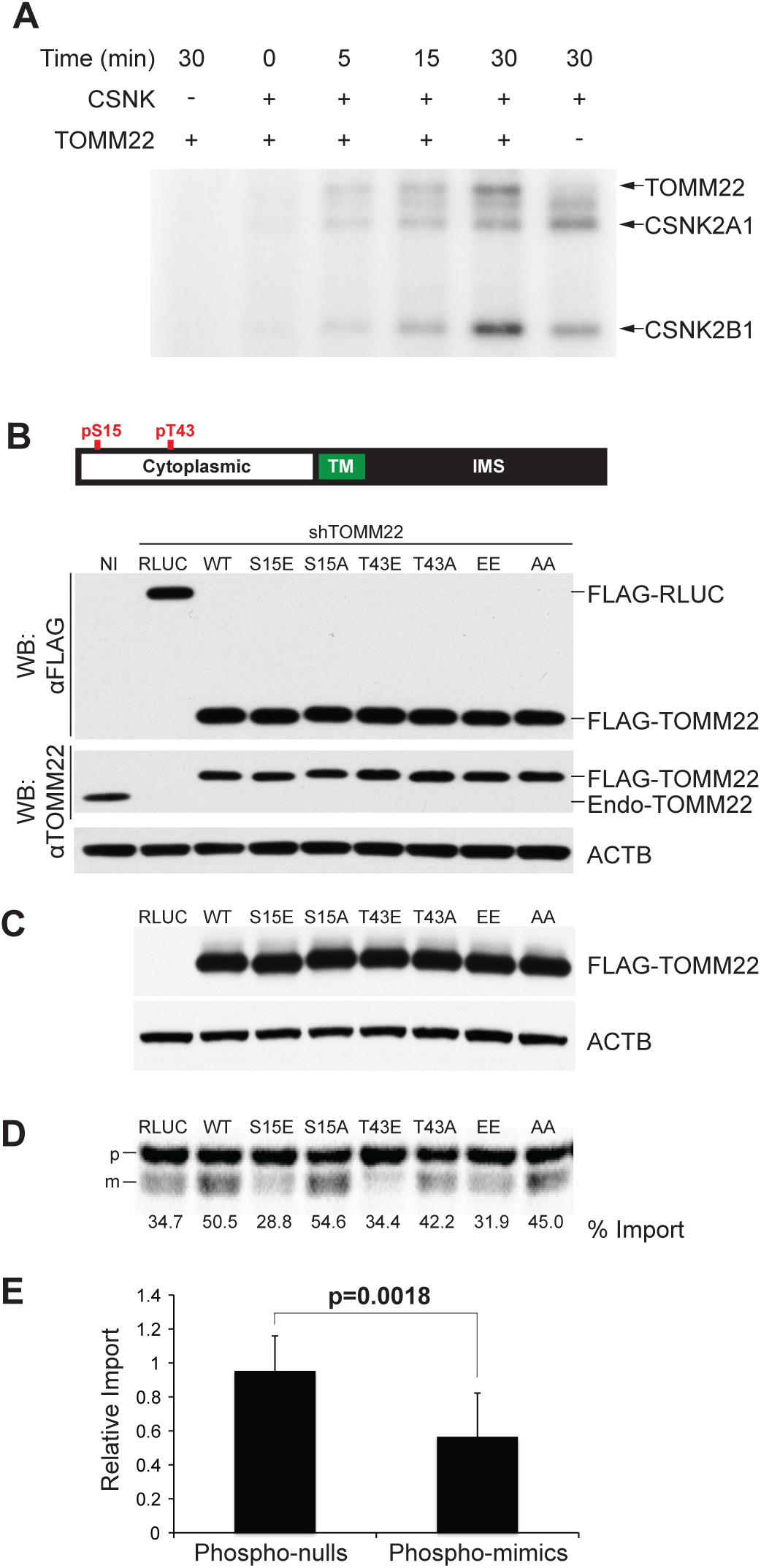
TOMM22 phosphorylation disrupts mitochondrial protein import. (A) TOMM22 phosphorylation by protein kinase CK2 was carried out with GST-TOMM22 in the presence protein kinase CK2 for increasing amounts of time. Reactions lacking TOMM22 or protein kinase CK2 were included as controls. Proteins were resolved by SDS-PAGE and visualized by autoradiography. (B) Schematic of TOMM22 protein and location of predicted protein kinase CK2 phospho-sites. Western blot analysis of whole cell lysates showing the relative expression of the N-terminally FLAG-tagged *Renilla luciferase* (36 kDa) and TOMM22 WT and mutant constructs (16.5 kDa) relative to endogenous TOMM22 (15.5 kDa) protein levels. (C) Western blot analysis of TOMM22 mutant expression from mitochondrial preparations. (D) Radiography of OCT import into mitochondria. Values represent the average from three independent experiments. p, precursor OCT; m, mature OCT. Data displayed are representative of three biological replicates (independent infections and mitochondrial isolations). (E) Bar graph showing the average relative import of OCT. Phospho-nulls and phospho-mimics have been grouped to display the effect of phosphorylation on import.

Given that TOMM22 is critical for protein import into mitochondria, we sought to assess if TOMM22 phosphorylation affected this process. In yeast, it has recently been shown that the phosphorylation of Tom22 by CK2 increased Tom22 localization to the mitochondrial membrane, and stimulated TOM complex formation, resulting in increased import of TOM complex members into the OMM (Schmidt *et al*. 2011). The *S. cerevisiae* and human TOMM22 proteins have 32% sequence similarity (Yano *et al*. 2000), and their functions are highly conserved, suggesting together with our screen data, that the functional consequences of this interaction may also be conserved in human cells. To investigate if phosphorylation of human TOMM22 was affecting the import of proteins into human mitochondria, a series of TOMM22 mutants were generated where the reported phospho-sites in the cytoplasmic domain of TOMM22 (serine 15 and threonine 43; Figure 5B) were each individually mutated to alanine (to block phosphorylation) or glutamic acid (to mimic phosphorylation), creating the clones S15E, S15A, T43E, and T43A. Additionally, the double phospho-null (AA) and double phospho-mimic (EE) were also generated. These clones also possessed an N-terminal FLAG tag so that exogenous TOMM22 could be distinguished from the endogenous protein by Western blot analysis. Wild-type TOMM22 and a *Renilla luciferase* (RLUC) construct were also N-terminally FLAG-tagged and used as controls. These sequences were stably expressed using a lentiviral system in HEK293T cells. To avoid interference from endogenous TOMM22, endogenous TOMM22 was depleted in cell lines stably expressing different TOMM22 phospho-mutants with an shRNA targeting the 3’UTR of endogenous TOMM22 (Figure 5B). To determine if TOMM22 phosphorylation influences the process of mitochondrial protein import, *in vitro* mitochondrial import assays were performed to assess the import capacity of mitochondria expressing mutant versions of TOMM22. Importantly, mitochondrial isolations from the mutant cell lines, and subsequent Western blot analysis of the relative levels of TOMM22-mutant protein, revealed that the localization of mutant TOMM22 to mitochondria was not impaired (Figure 5C). This was in contrast to the yeast system, where phospho-null mutants were impaired in their ability to localize to mitochondria (Schmidt *et al*. 2011).

To test the import capacity of mitochondria expressing mutant TOMM22, we performed *in vitro* mitochondrial import assays, which assess the ability of isolated mitochondria to import an *in vitro* transcribed and translated precursor protein, pre-ornithine transcarbamylase (pOTC), into the mitochondrial matrix (Terada *et al*. 1996). If imported into mitochondria, the pre-sequence of the OCT protein is cleaved off, generating a smaller protein, which can be distinguished as a lower molecular weight species by SDS-PAGE. Thus, protein import capacity can be estimated by quantifying the amount of mature OCT relative to the total amount in the reaction. Using mitochondria isolated from each of the stable cell lines described above (Figure 5B,C), it was found that the phospho-mimics were significantly impaired in the import of pOCT (Figure 5D). This is in contrast to the yeast report where phosphorylation increased import of TOM complex members and precursor proteins into mitochondria. These results indicate that the interaction between protein kinase CK2 and TOMM22 is conserved between yeast and man, but the human interaction appears to occur in a cell cycle dependent manner to decrease mitochondrial protein import into mitochondria during mitosis.

## DISCUSSION

We have described a new method, MaGiCaL-BiFC, for the sensitive and accurate detection of PPIs in mammalian cells. This system has numerous advantages over other PPI detection methodologies: (1) the lentiviral design of our system allows for the detection of PPIs in cell lines typically not amenable to high-throughput PPI screening approaches such as stem cells and terminally differentiated cell types, (2) the Gateway compatibility of our lentiviral vectors allows for the facile generation of genome-wide or cell-specific cDNA libraries using sequence-specific recombination, (3) the pooled-screening approach allows for the high-throughput analysis of PPIs in a cost-effective manner, precluding the requirement for abundant starting materials, (4) the ability to detect small proteins that may be missed by AP-MS techniques, (5) the irreversible folding of the fluorescent complex allows for the trapping and detection of transient interactions, or those with a weak but biologically important affinity, which other approaches would fail to detect, (6) the accurate study of native integral membrane proteins without altering localization signals, or rigorous purification schemes, and finally (7) the interaction can be localized, providing important information on the biological significance of the PPI. Importantly, MaGiCaL-BiFC provides an improvement over current mammalian BiFC methodologies, which although have been extremely valuable, have limitations with respect to the cell lines and proteins amenable to study.

We have applied MaGiCaL-BiFC to screen for TOMM22 protein interactions and have identified a list of highly enriched putative interaction partners of the TOM complex. The ease with which we were able to detect known and unknown interactors of TOMM22 highlights the applicability of this system to challenging proteins such as those with membrane-spanning domains. The specific identification of TOMM5 and TOMM6 in the N-terminal screens, and TOMM20 in the C-terminal screens, highlights the accuracy of this screening approach. Although not all of these putative interactors are likely *bone fide* interaction partners of TOMM22, the ability of BiFC to detect both binary and bystander protein interactions (e.g., in higher order complexes) (Vidi and Watts 2009) was reflected in the number of high confidence hits identified in these screens.

The ability of MaGiCaL-BiFC to detect the interaction between TOMM22 and protein kinase CK2 truly highlights the utility of this approach. Directed by previous phospho-proteomic data, we were able to confirm that this interaction is likely a mitotic-enriched event. This finding speaks to the effectiveness of MaGiCaL-BiFC at identifying transiently interacting proteins, which would otherwise be missed by more stringent technologies such as AP-MS.

The recent report that yeast CK2 interacts with and phosphorylates yeast Tom22 prompted us to pursue this interaction for follow-up (Schmidt *et al*. 2011). Interestingly, while the interaction is conserved in human cells, the consequences of this interaction are not. Importantly, in yeast, it was found that Tom22 is constitutively phosphorylated by CK2, and removal of this modification prevented mitochondrial protein import. This resulted from decreased association of un-phosphorylated Tom22 at the outer mitochondrial membrane. In contrast, recent reports using human cell lysates synchronized at various phases of the cell cycle suggest that unlike yeast, human TOMM22 is only phosphorylated during mitosis (Dephoure *et al*. 2008; Olsen *et al*. 2010). In support of this, the interaction between endogenous human protein kinase CK2 and TOMM22 was highly enriched in cells arrested in M-phase. Additionally, although the TOMM22 phosphorylation mutants did not show any differences in their ability to localize to mitochondria, we found that mutants mimicking the phosphorylation of either serine 15 or threonine 43 significantly blocked mitochondrial protein import. This is in direct contrast with the findings in yeast, but in line with the *Olsen et al.* study, which reported through enrichment analysis of their mitotic-specific phosphorylation events, that such events on metabolic proteins would likely decrease their activity (Olsen *et al*. 2010).

Given the role of protein kinase CK2 in cancer progression, it is logical to hypothesize that constitutive phosphorylation of TOMM22 may contribute to the metabolic changes commonly observed in human cell cancers. Protein kinase CK2 over-expression would likely result in constitutive phosphorylation of TOMM22 in cancer cells, decreasing mitochondrial protein import, and resulting in less functional mitochondria. This would be consistent with reports that have suggested that cancer cell mitochondria are generally smaller in size (Rehman *et al*. 2012). Moreover, decreased mitochondrial import of proteins important for respiration could contribute to the increased rates of glycolysis typical of cancer cells, or recent discoveries linking decreased mitochondrial respiration and cellular reprogramming (Xu *et al*. 2013). These ideas represent interesting avenues for future follow-up.

Altogether, this study has highlighted the utility of MaGiCaL-BiFC to identify both strong and weak or transient protein-protein interactions in mammalian cells, providing strong incentive for its application to other proteins important for normal and disease biology. This technology is particularly amenable to the discovery of membrane-based protein interactions, including proteins with membrane-spanning domains, in their native cellular environment. Thorough follow-up provided physical and functional validation of a number of the hits identified in this proof-of-concept screen, and validation of the interaction between TOMM22 and protein kinase CK2 has validated the utility of this approach to uncover novel biology that other protein-protein interaction methodologies have failed to uncover. When combined with other methodologies including Y2H, AP-MS and current PCA methodologies, MaGiCaL-BiFC provides an additional layer of coverage that can be utilized in the effort to map the functions of human proteins.

## ACKNOWLEDGEMENTS

We thank members of the Moffat laboratory for helpful discussions and I. Stagljar for comments on the manuscript. K.B. was supported by an NSERC graduate scholarship. This work was supported by funds from the Canadian Institutes for Health Research, Canadian Foundation for Innovation, Ontario Ministry of Research and Innovation and the Canadian Institute for Advanced Research to J.M. J.M. is a Tier II Canada Research Chair in Functional Genomics and Research Fellow at the Canadian Institute for Advanced Research.

