## Supplemental Figures for "Cell cycle regulation of mitochondrial protein import revealed by genome-scale pooled bimolecular fluorescence complementation screening"

*Donnelly Centre and Banting & Best Department of Medical Research, §Department of Molecular Genetics, University of Toronto, 160 College Street Room 802, Toronto, Canada, M5S3E1; †Muscle Health Research Centre, York University, Farquharson Life Sciences Building Room 302, 4700 Keele Street, Toronto, Canada, M3J1P3; ‡Samuel Lunenfeld Research Institute, 600 University Avenue, Toronto, Canada, M5G1X5

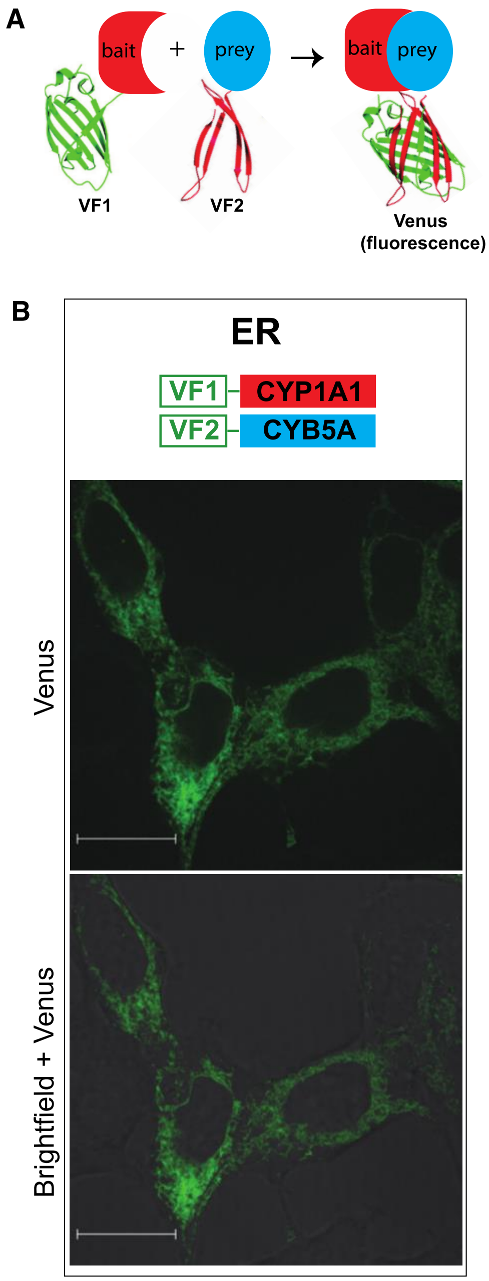


**Figure S1** MaGiCaL-BiFC assay with the endoplasmic reticulum associated proteins CYP1A1 and CYB5A. (A) Schematic representation of the bi-fluorescence complementation assay. (B) Results of a MaGiCaL-BiFC assay in HEK293T cells for two interacting proteins resident in the endoplasmic reticulum including CYP1A1 and CYB5A. Scale bar denotes 16 µm.

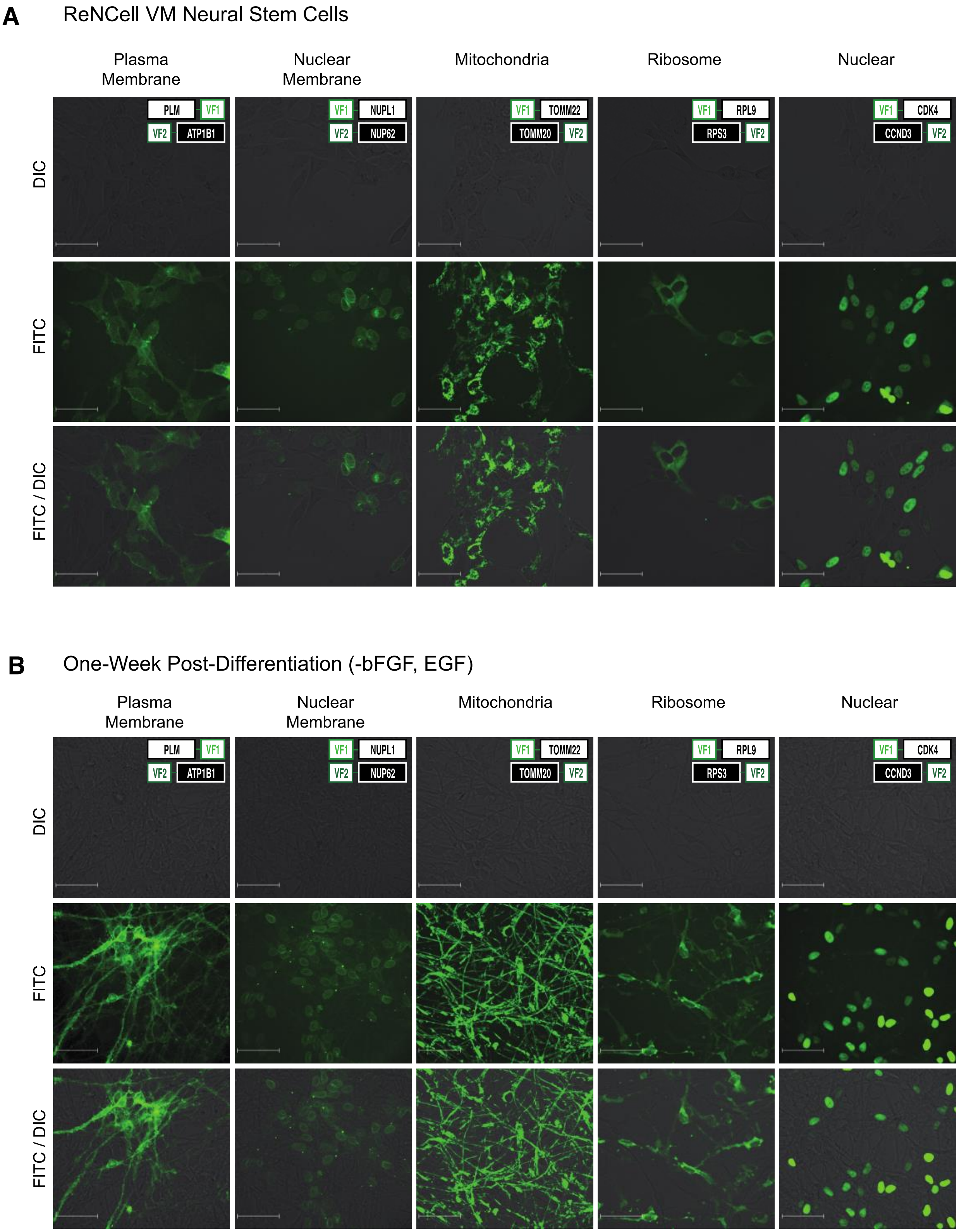
A
ReNCell VM Neural Stem Cells
Plasma Membrane
Nuclear Membrane
Mitochondria
Ribosome
Nuclear
DIC
FITC
FITC / DIC
PLM
VF1
VF2
ATP1B1
NUPL1
NUP62
TOMM22
TOMM20
RPL9
RPS3
CDK4
CCND3
B
One-Week Post-Differentiation (-bFGF, EGF)
Plasma Membrane
Nuclear Membrane
Mitochondria
Ribosome
Nuclear
DIC
FITC
FITC / DIC

**Figure S2** Pair-wise localization tests in neural stem cells and differentiated neurons. Expression clones for organelle-specific protein-protein interactions were generated for the plasma membrane, ribosome, mitochondria, nuclear membrane, and nucleus by LR site-specific recombination, with one protein of the pair introduced into both orientations of the bait vectors, and the other into both orientations of the prey vectors. (A) Lentivirus was generated and used to infect ReNCell VM neural precursors, either singly, as pairs (testing all four possible combinations) or with control constructs. Four-days post infection cells were imaged for Venus expression. (B) Infected cells were subsequently differentiated in the same media minus bFGF and EGF to induce neuronal differentiation. Media was replaced every two days for a total of 7 days before the differentiated cells were re-imaged for Venus expression.

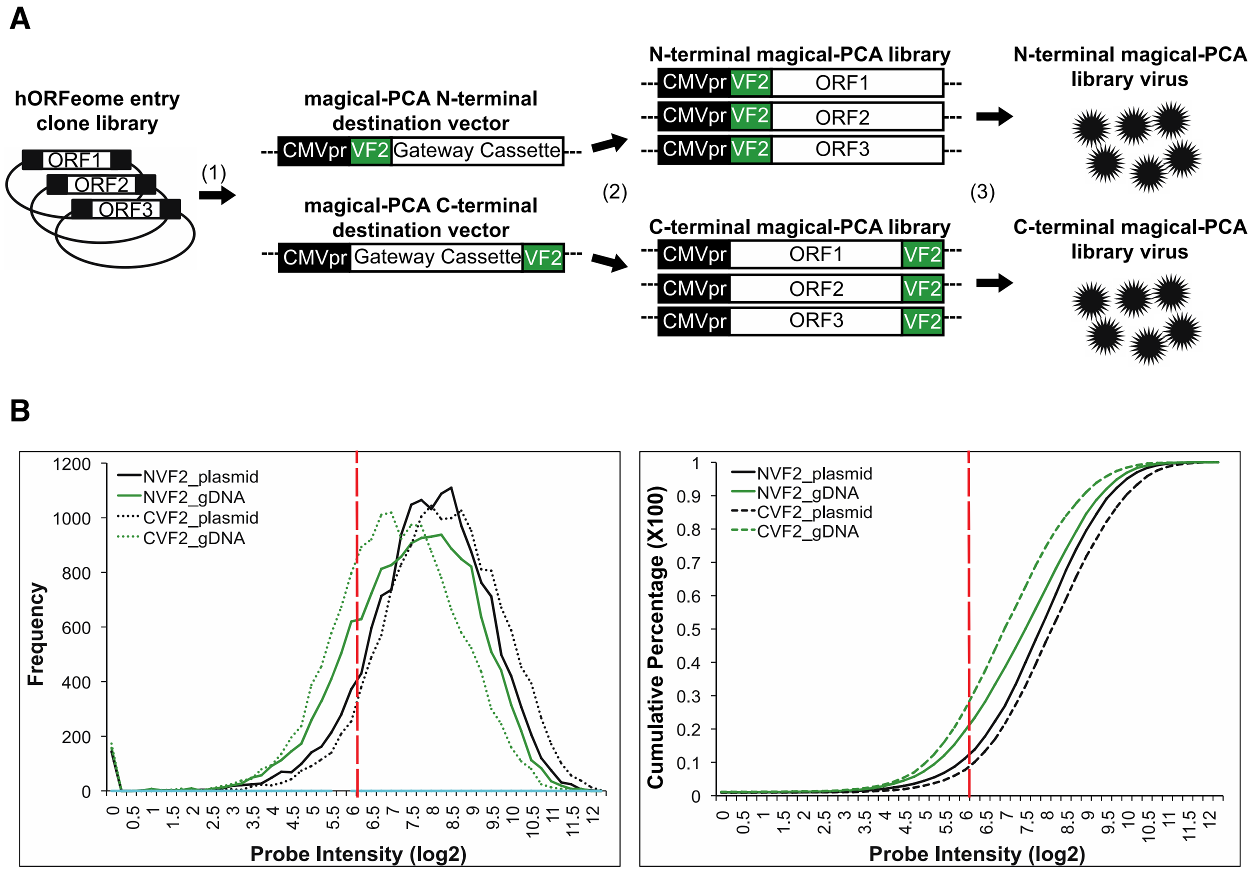


**Figure S3** Generation of MaGiCaL-BiFC human ORFeome v5.1 libraries. (A) (1) The hORFeome v5.1 entry clone library was transferred into both the N- and C-terminal MaGiCaL-BiFC destination vectors by LR site-specific recombination. (2) Transformants from each LR reaction were pooled together to generate separate N- and C-terminal MaGiCaL-BiFC libraries. (3) N- and C-terminal MaGiCaL-BiFC libraries were subsequently used to generate lentivirus for pooled infections. (B) ORF representation was estimated by hybridizing ORF amplicons from both the plasmid pools and genomic DNA from library transduced cells onto Affymetrix Gene 1.0 ST arrays. The dashed red line indicates background control signals.

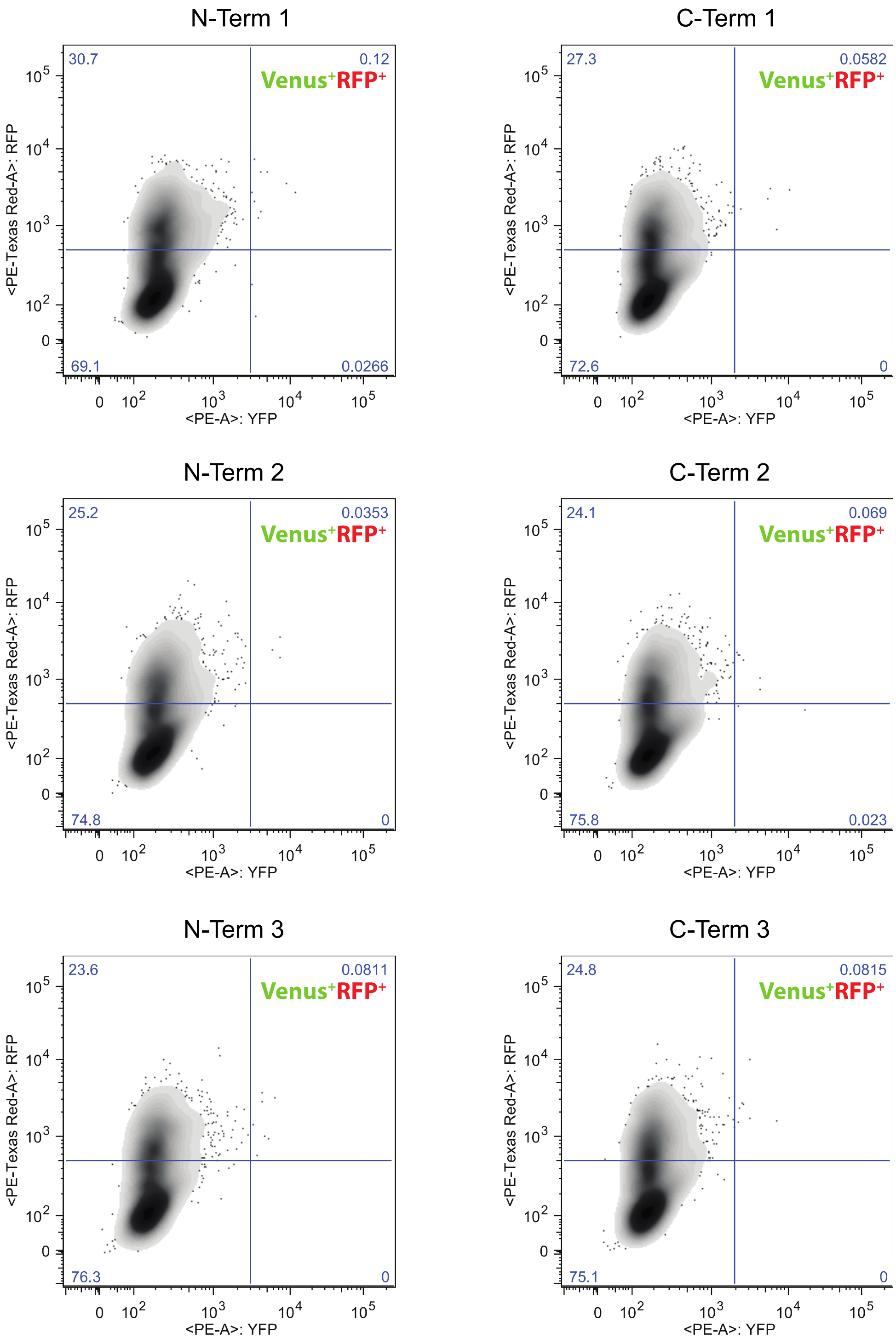


**Figure S4** FACS distributions for each of the N- and C-terminal TOMM22 pooled screens. Each of the 3 replicates for the N- and C-terminal library sorts is shown.

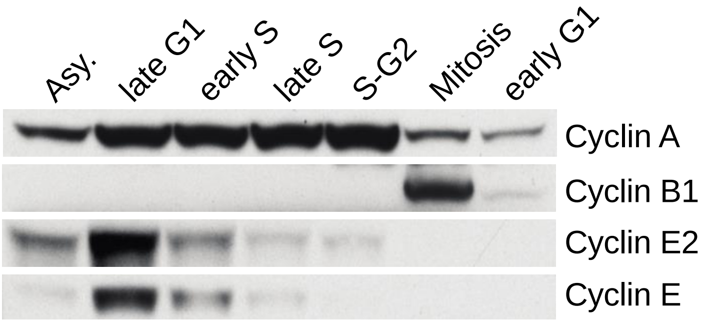


**Figure S5** Validation of cell cycle block/release samples. Western blot confirmation of successful cell cycle block/release samples using anti-cyclin antibodies specific to different phases of the cell cycle including Cyclin A (G1/S/G2), Cyclin B1 (mitosis), Cyclin E2 and Cyclin E (late G1/early S-phase).
